# Functional Redundancy of ZmSWEET6a/b in Mediating Sugar Transport and Redox Homeostasis for Maize Primexine Formation

**DOI:** 10.64898/2026.01.25.701620

**Authors:** Yan Zhang, Shuangtian Bi, Fengkun Sun, Jiajun Bu, Yurong Wang, Mateus Mondin, Zhaobin Dong, Weiwei Jin, Wei Huang

## Abstract

Male gametophyte development in maize requires precise coordination of carbohydrate allocation, yet the molecular mechanisms linking sugar transport to early pollen wall formation remain poorly understood. Although sugar transporters, particularly SWEET family members, are known to regulate male fertility in *Arabidopsis* and rice, their roles in maize microsporogenesis—especially in primexine assembly and redox homeostasis—have not been functionally characterized. Here, we identify two anther-specific Clade II SWEET genes, *ZmSWEET6a* and *ZmSWEET6b*, encoding redundant plasma membrane hexose transporters. Loss of both genes leads to complete male sterility, accompanied by disrupted deposition of primexine polysaccharides (pectin and xylan), premature accumulation of reactive oxygen species (ROS), and ectopic programmed cell death (PCD) in the tapetum. We demonstrate that *ZmSWEET6a/b*-mediated hexose transport is essential for maintaining sugar homeostasis and redox balance during early microsporogenesis. These findings establish a direct mechanistic link between sugar transport, ROS signaling, and pollen wall development in maize, providing new insights into the integration of metabolic and developmental pathways during plant reproduction.

## 1. Introduction

Male gametophyte development is a tightly regulated process, and disruptions at critical stages often lead to male sterility ^[1]^. The tapetum—the innermost somatic layer of the anther wall—plays a central role by supplying nutrients, enzymes, and structural precursors to developing microspores ^[2, 3]^. During meiosis, microspores become symplastically isolated by a callose wall, making the tapetum their essential source of support ^[2]^. A key tapetal function is facilitating the formation of the pollen exine, which begins with the establishment of a polysaccharide-rich primexine template ^[3, 4]^. Notably, reactive oxygen species (ROS) homeostasis is crucial during this process, as excessive ROS accumulation can trigger premature tapetal programmed cell death (PCD) and disrupt exine patterning ^[5]^. In maize, several mutants such as *Zmenr1, Zmms33* and *atp6c* exhibit early ROS bursts accompanied by defective pollen wall formation ^[6-8]^, underscoring the importance of redox balance in coordinating tapetal function and exine development.

The pollen exine, a resistant outer wall essential for viability and fertilization, consists of a sculptured sexine and a smooth nexine ^[3]^. Its formation begins when microspores are encased in a temporary callose wall ^[9]^. A cellulose-pectin matrix, the primexine, is then deposited onto the microspore surface, serving as the spatial template that guides the placement of sporopollenin precursors supplied by the tapetum ^[10, 11]^. Subsequent sporopollenin deposition builds the definitive exine layers ^[12]^. Primexine defects directly lead to pollen abortion, as demonstrated in *Arabidopsis* mutants such as *arf17, npu*, and *rpg1* ^[13-15]^, and in rice mutants including *tms10* and *dmd1* ^[16, 17]^. In maize, while several male-sterile mutants with exine abnormalities have been identified—affecting lipid metabolism (*ipe1, ms6021*) or transcriptional regulation (*ms1, ms7*) —their direct connection to primexine biogenesis remains unclear^[18, 19]^. An exception is *ms39*, where disrupted callose synthesis secondarily abolishes primexine formation^[20]^. Overall, the molecular mechanisms underlying primexine assembly in maize are still poorly understood.

As the development of photosynthetically inactive and symplastically isolated microspores entirely depends on external sugar supply, the precise allocation of carbohydrates mediated by sugar transporters (STs) is fundamental for male fertility ^[2, 21]^. In both *Arabidopsis* and rice, mutations in key ST genes, including sucrose transporters (*AtSUC1, OsSUT1*) ^[22, 23]^, SWEET family exporters (*AtSWEET8/RPG1, AtSWEET13/RPG2, OsSWEET11a/11b*) ^[15, 24, 25]^, and monosaccharide/H□ symporters (*AtSTP2, OsMST7/8*) ^[26, 27]^—commonly lead to male sterility, with defects spanning pollen maturation, germination, and anther dehiscence. While the roles of STs in these later developmental stages are relatively well-documented, their specific functions and regulatory networks during the critical early stages, including meiosis, tetrad release, and primexine scaffold formation, remain poorly understood. Notably, *AtSWEET8/RPG1* is the only ST directly linked to primexine assembly ^[15]^, highlighting a key gap in understanding early sugar-mediated regulation of exine development.

In maize, while members of the SWEET family have been implicated in critical agronomic processes such as phloem loading (*ZmSWEET13a, b, c*), seed filling (*ZmSWEET4c*), and likely source-sink allocation, their specific roles in male reproductive development, particularly during early microsporogenesis ^[28, 29]^, remain virtually unexplored. Recent phylogenetic and transcriptomic analyses have identified several anther-preferential or anther-specific SWEET genes in maize, suggesting their potential involvement in male fertility. Here, we characterize *ZmSWEET6a* and *ZmSWEET6b*, two anther-specific Clade II SWEET genes encoding redundant plasma membrane hexose transporters. We demonstrate that their loss disrupts primexine polysaccharide (pectin/xylan) deposition, triggers premature ROS bursts, induces ectopic tapetal PCD, and ultimately causes male sterility. This work establishes a direct mechanistic link between hexose transport, redox balance, and pollen wall development in maize, offering new insights into maize microsporogenesis.

## 2. Materials and methods

### 2.1. Plant materials and growth conditions

The maize (*Zea mays* ssp. *mays* L.) *Zmsweet6a, Zmsweet6b* single mutants and the *Zmsweet6a/6b* double mutants were generated in the LH244 inbred line by CRISPR-Cas9 targeting ^[30]^. Three independent mutant lines for each gene were obtained: *Zmsweet6a*^*d2*^, *Zmsweet6a*^*i1d16*^ and *Zmsweet6a*^*d16*^ for *Zmsweet6a*; and *Zmsweet6b*^*d2*^, *Zmsweet6b*^*i1d7*^, and *Zmsweet6b*^*d13*^ for *Zmsweet6b*. Each line carries a deletion in the third exon of the corresponding gene. The *Zmsweet6a/6b* double mutant lines were subsequently generated by crossing the respective single mutants. Unless otherwise specified, all plants, including the WT LH244 and the mutants, were cultivated during the summer at the experimental farm of China Agricultural University (CAU; 40°01′N, 116°16′E). During the winter, plants were grown in Sanya, Hainan Province.

### 2.2. Plasmid construction and plant transformation

For targeted mutagenesis of *ZmSWEET6a*, a CRISPR-Cas9 plasmid was constructed by cloning guide RNA(s) targeting a 19-bp spacer sequence within the *ZmSWEET6a* coding region into the binary vector pBUE411 ^[30]^. *ZmSWEET6b* and the *ZmSWEET6a/6b* double mutant were obtained by screening the off-target events. Transgenic plants were identified by genomic PCR using gene-specific primers, followed by sequencing verification. All primers used for plasmid construction are listed in Supplementary Table 3.

### 2.3. Phenotype observation

Morphological characteristics of whole maize plants and maize tassels were photographed using a Nikon D7100 digital camera (Nikon, Tokyo, Japan). Images of anthers were acquired using an Olympus SZX10 stereomicroscope (Olympus, Tokyo, Japan). Micrographs of pollen grains stained with a 1% I□–KI solution were captured using an Olympus CX41F microscope (Olympus, Tokyo, Japan).

### 2.4. Sequence alignment and phylogenetic analysis

To identify orthologs and paralogs of ZmSWEET6a, BLAST searches were performed using its protein sequence against the following databases: Ensembl Plants (https://plants.ensembl.org/index.html), the National Center for Biotechnology Information (https://www.ncbi.nlm.nih.gov/), Phytozome (https://phytozome.jgi.doe.gov/pz/portal.html), and the maize genome database (https://www.maizegdb.org/). Multiple sequence alignment of protein sequences encoded by *SWEET* genes from *Arabidopsis thaliana, Oryza sativa*, and *Zea mays* was conducted using the ClustalW algorithm implemented in the msa R package. The resulting alignment was subsequently converted into a format suitable for phylogenetic analysis (phyDat) using the phangorn package. Based on this, a Maximum Likelihood (ML) phylogenetic tree was constructed employing the Jones-Taylor-Thornton (JTT) amino acid substitution model. The robustness of the phylogenetic tree topology was assessed with 1,000 bootstrap replicates. Four major evolutionary clades (Clade I–IV) were manually delineated based on branching features, and representative genes were designated for each.

### 2.5. Real-time Quantitative PCR analysis

To determine the relative expression levels of *Zmsweet6a* and *Zmsweet6b* in the vegetative organs (root, leaf, stem, ear) of maize plants and in anthers at different developmental stages, real time quantitative PCR (RT-qPCR) was performed. Total RNA extraction, reverse transcription, and RT-qPCR assays were carried out following previously described protocols ^[31]^. The primer sequences used for quantitative PCR are listed in Supplementary Table 3.

### 2.6. Messenger RNA *in situ* hybridization

The meiotic-staged florets of LH244 were fixed in 3.7% formalacetic-alcohol (FAA) and *in situ* hybridization was followed the previously described protocol ^[32]^. Sense and antisense probes of *ZmSWEET6* were generated by PCR amplification with SP6 and T7 polymerase, respectively. The gene-specific primers are listed in Supplementary Table 3.

### 2.7. Immunofluorescence staining

Meiotic tassels from WT (LH244) and *Zmsweet6a/6b* plants were used for immunoblot analysis. The immunoblotting procedure was performed as previously described ^[33, 34]^. The primary antibodies JIM5 (CarboSource Service, Athens, GA, USA) and LM10 (Megazyme, Bray, Ireland) were used at a dilution of 1:100. A goat anti-rat immunoglobulin G antibody conjugated to horseradish peroxidase (ABclonal) was used as the secondary antibody at a dilution of 1:5000.

### 2.8. Observation of meiotic chromosomes and callose

Chromosome preparation during meiosis was performed as described previously ^[31]^. The chromosomes were stained with DAPI and toluidine blue, and chromosome images were captured under an Olympus BX61 fluorescence microscope. DAPI-stained samples were observed under a UV excitation filter (excitation wavelength: 350–360□nm, emission wavelength: 450–460□nm), whereas toluidine blue-stained samples were examined under bright-field illumination.

### 2.9. Cytological analysis and microscopy

Fresh anthers from WT (LH244) and *Zmsweet6a/6b* mutant plants, spanning developmental stages 5 to 12, were collected and processed separately. For semithin sectioning, anthers were fixed in FAA solution. For transmission electron microscopy (TEM) analysis, anthers were fixed in a mixed solution containing 4% paraformaldehyde and 5% glutaraldehyde, followed by post-fixation in 1% osmium tetroxide overnight. For semithin sectioning, fixed anthers were dehydrated through a graded ethanol series (30%, 50%, 70%, 80%, 95%, and 100%, v/v). The samples were then infiltrated and embedded using graded mixtures of Spurr’s resin (resin/propylene oxide ratios of 1:3, 1:1, and 3:1, v/v) and polymerized at 70°C for 14 hours. Sections of 2 μm thickness were cut using a Leica RM2265 rotary microtome, stained with 0.1% (w/v) crystal violet (Sigma-Aldrich) for 5 minutes, and observed under an Olympus BX-53 microscope. All subsequent procedures for TEM analysis were performed according to a previously described method ^[35]^.

### 2.10. Subcellular localization

The CDSs of *ZmSWEET6a* and *ZmSWEET6b* were cloned and inserted into the pCAMBIAsuper1300 by homologous recombination, resulting in the constructs *ZmSWEET6a*-GFP and *ZmSWEET6b*-GFP, respectively. The constructs were transformed into maize protoplasts and *Nicotiana benthamiana* leaves, and GFP fluorescence in protoplasts was visualized under a confocal scanner microscope (LSM 900, Zeiss). The pCAMBIAsuper1300 vector containing the green fluorescent protein (GFP) alone served as a negative control. Primers used for plasmid construction are listed in Supplementary Table 3.

### 2.11. Complementation assays of ZmSWEET6a and ZmSWEET6b in the yeast

The ZmSWEET6a-PDR196 and ZmSWEET6b-PDR196 vectors were constructed and subsequently transformed into *Escherichia coli* for plasmid amplification. Following plasmid extraction, competent cells of the *EBY*.*VW4000* yeast strain were prepared. The recombinant plasmids were then transformed into the *EBY*.*VW4000* competent cells. After incubation at 30°C for 3 hours, 100 μL of the transformation mixture was spread onto SD-ura plates with maltose as the sole carbon source and incubated at 30°C for 1–3 days. Single colonies were picked and inoculated into 1 mL of YPD medium for shaken incubation at 30°C for 1–2 hours. Subsequently, 100 μL aliquots of the yeast cultures were transferred onto SD-ura plates containing either glucose or fructose as the sole carbon source. The plates were incubated at 30°C for 1–3 days, and colony growth was documented by photography using a Nikon D7100 digital camera (Nikon, Tokyo, Japan).

### 2.12. Sugar-targeted metabolomic analysis

Anthers of WT (LH244) and *Zmsweet6a/6b* double mutant at developmental stages 6 and 7 were collected and immediately frozen in liquid nitrogen for storage. The freeze-dried anther samples were ground using a mixer mill (MM 400, Retsch). Approximately 20 mg of the resulting powder was accurately weighed and extracted with 500 μL of a methanol: isopropanol: water mixture (3:3:2, v/v/v). The mixture was vortexed for 3 minutes and sonicated for 30 minutes, followed by centrifugation at 12,000 rpm for 3 minutes at 4°C. A 50 μL aliquot of the supernatant was combined with 20 μL of internal standard solution (1000 μg/mL) and dried under a stream of nitrogen gas. The dried samples were then transferred to a freeze-dryer. The resulting residue was used for subsequent derivatization. After derivatization, the reaction mixture was diluted to an appropriate concentration and analyzed by gas chromatography-mass spectrometry (GC-MS) based on the Agilent 8890-5977B platform. Data were processed and analyzed using GraphPad Prism software (version 8.0.2).

### 2.13. RNA sequencing and transcriptome analysis

Meiotic anthers of *Zmsweet6a/6b* and WT (LH244) were collected for RNA extraction and the subsequent RNA-seq, with three biological replicates for each genotype. RNA-seq and differentially expressed genes (DEGs) analysis are listed in Supplementary Table 4, 6. The Gene Ontology (GO) enrichment analysis of DEGs was performed with g:Profiler (https://biit.cs.ut.ee/gprofiler/gost). The corresponding GO enrichment terms and associated gene lists are provided in Supplementary Table 5.

### 2.14. NBT staining

Fresh anthers of WT (LH244) and *Zmsweet6a/6b* from stage 6 to stage 10 were immersed in 10 mM potassium-citrate buffer (pH 6.0) and vacuum-infiltrated for 15 min, then exchanged with 0.5 mM NBT solution and incubated for 3 h at room temperature. The stained anthers were washed with 70% (v/v) ethanol thrice and immersed in FAA solution (Coolabor, China) overnight. Then, anthers of WT and *Zmsweet6a/6b* were embedded in paraffin (Sigma, USA). The subsequent procedures were performed as described previously^[36]^. The signals were observed and photographed using an Olympus CX41F microscope (Olympus, Tokyo, Japan).

### 2.15. Measurement of ROS content assay

The collected fresh anthers of WT and *Zmsweet6a/6b* were washed using PBS for 5 min, then stained with H2DCF-DA (Sigma-Aldrich) according to established protocols ^[37]^. After washing thrice with PBS, fluorescence was observed under a confocal laser-scanning microscope (M205FA, Leica, Japan). The relative fluorescence intensities of ROS were quantified using the ImageJ software.

### 2.16. TUNEL assay

Anthers of WT and *Zmsweet6a/6b* embedded in paraffin (Sigma, USA) were used for TUNEL (Terminal deoxynucleotidyl transferase dUTP Nick-End Labeling) analysis. The subsequent procedures were performed as described previously^[36]^. Nick-end-labeled nuclear DNA fragments in the WT and *Zmsweet6a/6b* were visualized with a Dead End Fluorometric TUNEL Kit (DeadEndTM Fluorometric TUNEL System, Promega). TUNEL signals were observed and photographed using a fluorescence confocal scanner microscope (TCS-SP8, Leica).

## 3. Results

### *3*.*1. ZmSWEET6a* and *ZmSWEET6b* play redundant roles in maize male fertility

In our previous study, we found that the sugar metabolism gene *INVAN6* plays a critical role in ensuring the normal progression of meiosis under high-temperature conditions ^[38]^. We hypothesized that during maize microsporogenesis, additional sugar-related genes are functionally involved. By analyzing published transcriptome datasets ^[39, 40]^ and examining the expression of genes involved in sugar metabolism and transport in anthers, we identified two SWEET-encoding genes, ZmSWEET6a and ZmSWEET6b, that are highly expressed in anthers, particularly in meiocytes (Fig. S1A, B; Table S1). Phylogenetic analysis indicates that the products encoded by *ZmSWEET6a* and *ZmSWEET6b* have a 97.54% amino acid sequence identity, and both genes are predominantly expressed in meiotic tassel, suggesting functional redundancy (Fig. S1C, S2). Consistent with this hypothesis, our results show that neither the *Zmsweet6a* nor the *Zmsweet6b* single mutant exhibits phenotypic differences relative to the wild type (Fig. 1A, B and S3). However, simultaneous knockout of both *ZmSWEET6a* and *ZmSWEET6b* results in complete male sterility in the double mutant (Fig. 1B).

**Fig 1.**
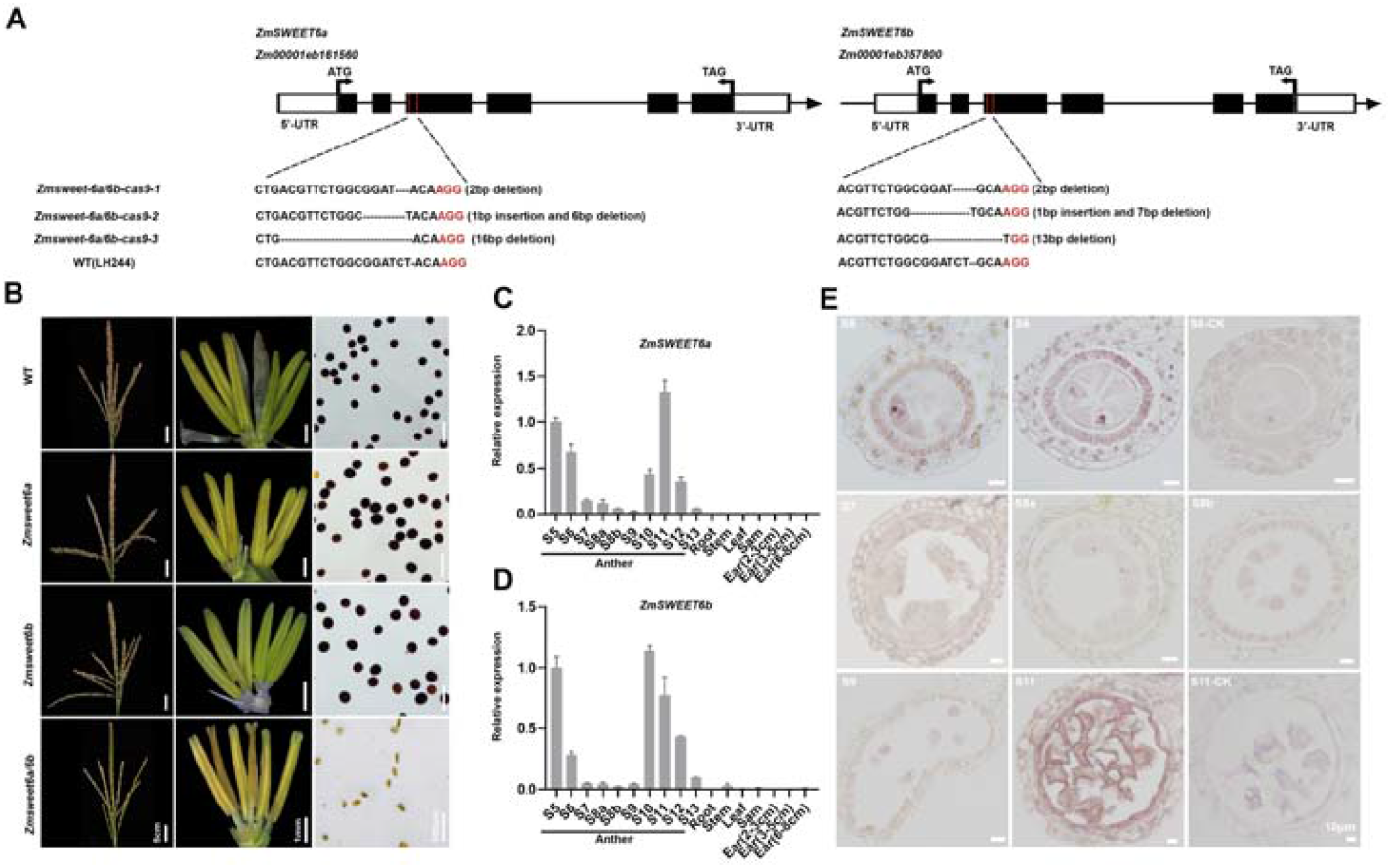
*ZmSWEET6a* and *ZmSWEET6b* are specifically expressed in anthers and essential for male fertility in maize. **(A)** Three CRISPR-Cas9 *Zmsweet6a/6b* double mutant lines. **(B)** Tassels, anthers, and pollen grains (stained by iodine-potassium iodide) of the WT (LH244), *Zmsweet6a* single mutant, *Zmsweet6b* single mutant and *Zmsweet6a/6b* double mutant. **(C D)** *ZmSWEET6a* and *ZmSWEET6b* are expressed specifically in maize anthers based on quantitative real-time PCR (qPCR) analysis. Anther developmental stages are indicated as S5–S12. Data are the mean ± SD of the relative expression values normalized to the internal control ZmUbi2 from three biological replicates. **(E)** *In situ* hybridization with mRNA probes was employed to detect *ZmSWEET6* gene signals in WT maize anthers during stages S5 to S11. In B-E, each experiment was repeated three times independently. Scale bars are shown in the images.

We next examined the expression patterns of *ZmSWEET6a* and *ZmSWEET6b* in detail. Both genes are predominantly expressed in anthers, consistent with the observation that the the *Zmsweet6a/6b* double mutant displays male sterility without affecting other agronomic traits (Fig. S1A, B, S3). Interestingly, RT-qPCR (Reverse Transcription Quantitative Real-time PCR) showed *ZmSWEET6a* and *ZmSWEET6*b exhibit highly coordinated dynamic expression, with pronounced upregulation during two key anther developmental windows: stages S5–S6 (meiosis) and S10–S12 (microspore development) (Fig. 1C, D). Furthermore, mRNA *in situ* hybridization showed that *ZmSWEET6a/6b* transcripts are primarily enriched in the tapetum and germ cells at stages 5–6 and stage 11 (Fig. 1E).

Collectively, based on their overlapping expression patterns and the sterile phenotype of the double mutant, we conclude that *ZmSWEET6a* and *ZmSWEET6b* play a crucial and functionally redundant roles in maize male gametophyte development.

### 3.2. The *Zmsweet6a/6b* double mutant shows premature tapetum vacuolization and aberrant pollen exine formation

Both *ZmSWEET6a* and *ZmSWEET6b* are highly expressed in the tapetum and pollen mother cells (PMCs) during early meiosis. We initially examined meiosis in PMCs of the *Zmsweet6a Zmsweet6b* double mutant but observed no significant abnormalities (Fig. S4). This prompted us to investigate whether the male sterility phenotype arises from defects in tapetal development. To elucidate the cause of pollen sterility in the *Zmsweet6a/6b* double mutant, we performed a comprehensive histological analysis of transverse anther sections from wild-type (WT) and mutant plants across developmental stages 5 to 12 (Fig. 2).

**Fig 2.**
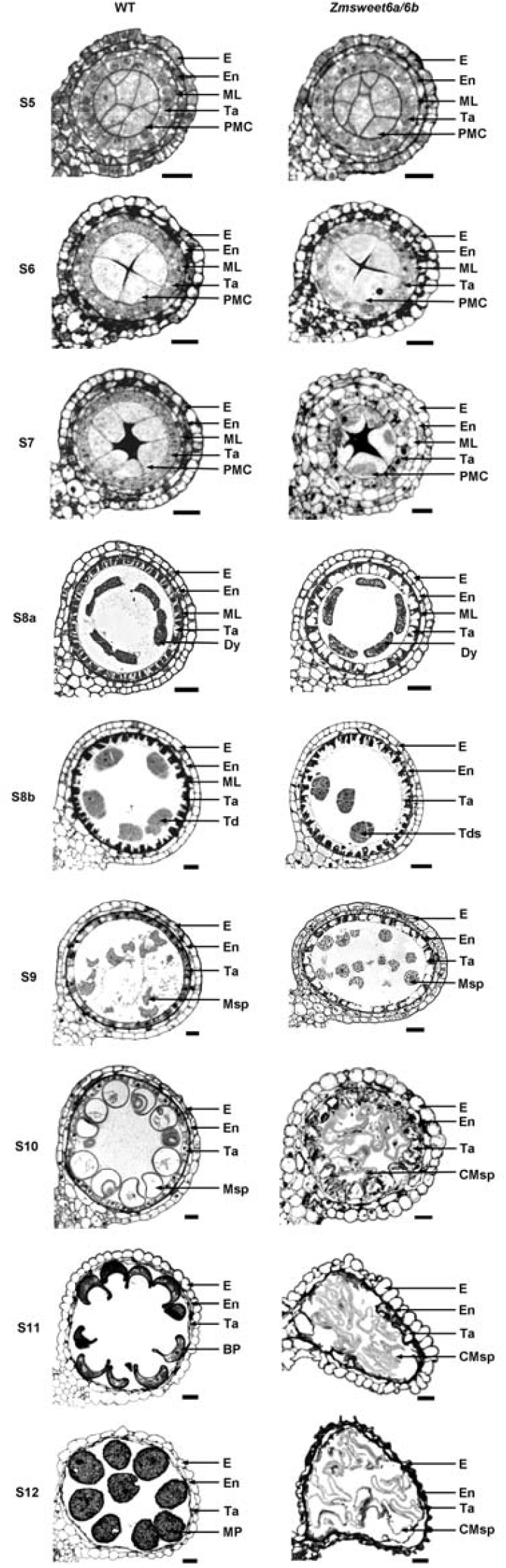
Comparative analysis of anther and pollen development in WT and *Zmsweet6a/6b* double mutant using transverse sections. E, epidermis, En, inner wall of locule, ML, middle layer; Ta, tapetum, PMC, pollen mother cell, MC meiotic cell, Dy, dyad, Td, tetrad, Msp, microspore CMsp, collapsed microspore, BP, binucleate pollen, MP, mature pollen. Bar, 20 μm.

At stage 5, anthers of both WT and *Zmsweet6a/6b* plants appeared morphologically indistinguishable (Fig. 2). However, prominent abnormalities emerged in the mutant beginning at stage 6. In *Zmsweet6a/6b*, obvious vacuolization was already evident in the tapetum at stage 6 and intensified progressively during subsequent stages. In contrast, tapetal vacuolization in the WT did not commence until stage 8a (Fig. 2). Although germ cells within the anther locules underwent two rounds of meiosis to form uninucleate microspores, a process that proceeded normally in both genotypes, subsequent pollen development diverged markedly. By stage 10 in the WT (Fig. 2), microspores became vacuolated, enlarged, and adopted a rounded or crescent shape with uniformly thickened pollen walls. In contrast, *Zmsweet6a/6b* microspores failed to undergo vacuolation, exhibited irregular shapes, and displayed abnormally thick and unevenly deposited pollen walls. By stage 12 (Fig. 2), WT pollen grains regained a spherical morphology following a transient sickle-shaped phase at stage 11, whereas *Zmsweet6a/6b* pollen grains remained misshapen due to a failure to accumulate starch.

To further dissect the developmental defects in the double mutant, we compared anther ultrastructure between WT and *Zmsweet6a/6b* plants from stages 5 to 10 using transmission electron microscopy (TEM) (Fig. 3, S5). Consistent with semi-thin section observations, TEM revealed that anther structure was normal in *Zmsweet6a/6b* at stage 5. However, by stage 6, the tapetum in the mutant already exhibited pronounced vacuolization (Fig. 3A, B), which became significantly more severe than in the WT throughout stages 7–9 (Fig. 3C–F). As development proceeds, small, spherical Ubisch bodies form on the surface of the tapetum and deliver sporopollenin to microspores to support pollen wall formation ^[41]^. Although the tapetum in the *Zmsweet6a/6b* mutant already exhibits abnormalities as early as stage 6, Ubisch bodies are still formed but appear to be even more densely distributed (Fig. 3F, G).

**Fig 3.**
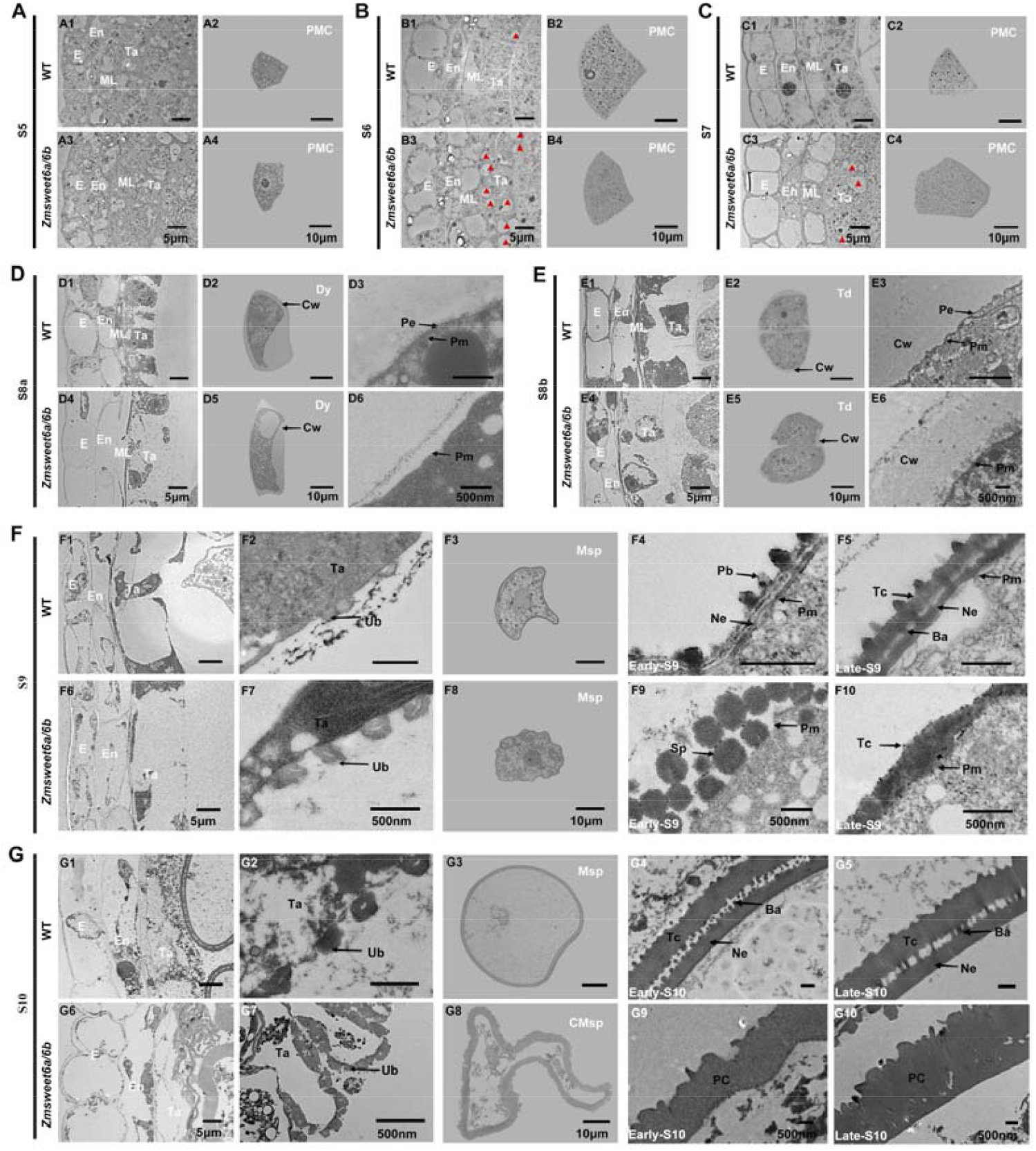
Transmission electron microscopy-based (TEM) examination of anther wall structures, microspores, Ubisch bodies, and exine formation in WT and *Zmsweet6a/6b*. **(A-C)** Four anther wall layers and pollen mother cells in WT and *Zmsweet6a/6b* double mutant at stages S5, S6, and S7. E, epidermis, En endothecium, ML middle layer; Ta tapetum, PMC pollen mother cell. **(D-E)** The structure of anther wall four layers and anther primary wall in WT and *Zmsweet6a/6b* double mutant at stages S8a and S8b. Dy, dyad, Td, tetrad, Cw, callose wall, Pe, primexine, Pm, plasma membrane. **(F-G)** Morphological characteristics of anther wall layers, Ubisch body, pollen mother cells, and anther outer wall in WT and *Zmsweet6a/6b* double mutant at stages S9 and S10. Ub, Ubisch body, Ta, tapetum, Msp, microspore, CMsp, collapsed microspore, Pb, probaculae; In, intine, Ne, nexine; Tc, tectum, Sp, sporopollenin, Ba, baculua, Pc, pollen coat. Scale bars are shown in the images.

In addition to premature tapetal vacuolization, pollen wall development was severely disrupted in *Zmsweet6a/6b*. From stages 5 to 8b, PMCs undergo meiosis and are surrounded by a callose wall composed of β-1,3-glucan. Toluidine blue staining confirmed that callose deposition occurred normally in both WT and *Zmsweet6a/6b* (Fig. S4B–J), and TEM further verified the presence of intact callose walls around dyads and tetrads in both genotypes (Fig. 3D, E). Following meiosis, the callose wall need to be degraded to release haploid microspores ^[42]^. Prior to this degradation, WT microspores typically develop a thin primexine layer between the plasma membrane and the callose wall, accompanied by characteristic undulations of the plasma membrane (Fig. 3D3, E3). Nevertheless, neither the undulation of plasma membrane nor the primexine formation are happened in *Zmsweet6a/6b* (Fig. 3D6, E6).

Plasma membrane undulation is proposed to direct the spatial arrangement of baculae, with their precursors, probaculae, specifically forming at the crests of the undulating membrane ^[43, 44]^. In WT microspores at early stage 9, sporopollenin delivered by the tapetum initiates the formation of probaculae and nexine. By late stage 9, a mature exine architecture comprising the tectum, bacula, and nexine is established (Fig. 3F4, F5). However, *Zmsweet6a/6b* failed to form either probaculae or nexine. Instead, sporopollenin accumulated as spherical aggregates on the microspore surface, eventually forming a dense, amorphous pollen coat (Fig. 3F9, F10). At stage 10, the WT exine further thickened but retained a uniform bilayer structure consisting of tectum and nexine with regularly spaced bacula (Fig. 3G4, G5). In the mutant, however, the microspore wall also thickened but exhibited extreme heterogeneity in thickness and lacked organized exine layers (Fig. 3G9, G10).

Collectively, these findings demonstrate that *ZmSWEET6a* and *ZmSWEET6b* are essential for maintaining tapetal integrity and orchestrating proper exine patterning. Their loss in *Zmsweet6a/6b* triggers premature tapetal vacuolization, disrupts primexine formation and sporopollenin deposition, and ultimately leads to complete male sterility in maize.

### 3.3. Loss of ZmSWEET6a/6b disrupts pectin and xylan deposition during primexine assembly

Previous studies have established xylan and pectin (derived from these precursors) as core components of the *Arabidopsis* primexine ^[4, 45]^. In this study, we observed a complete failure of primexine formation in *Zmsweet6a/6b* mutants. We therefore asked whether this defect is linked to impaired biosynthesis or deposition of pectin and xylan. To address this question, we further performed immunofluorescence staining on anther sections from WT and mutant plants at stages 5 to 10 (Fig. 4). Staining with the JIM5 antibody, which recognizes partially methyl-esterified HG ^[34]^, revealed a dense, uniform fluorescent ring surrounding WT microspores at stages 6 and 7. This indicates the continuous and robust deposition of the pectin matrix supporting the primexine formation. In contrast, *Zmsweet6a/6b* microspores displayed markedly weaker and fragmented JIM5 signals at the same stages (Fig. 4). The absence of a well-defined ring-like pattern indicates a disruption in the assembly of a structurally intact pectin scaffold in the double mutant.

**Fig 4.**
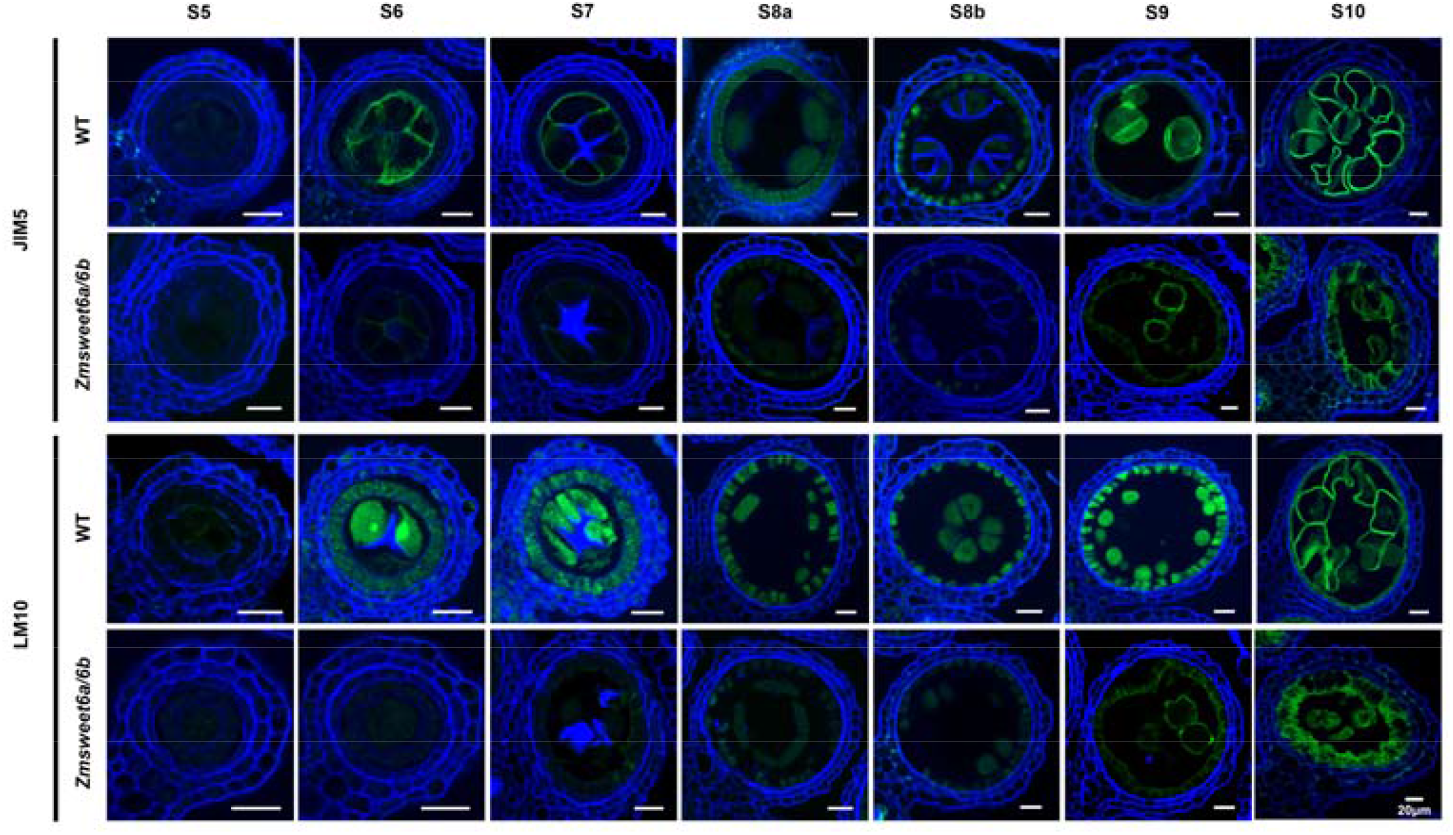
Mutations in *ZmSWEET6a* and *ZmSWEET6b* disrupt the normal accumulation of pectin and Xylan during pollen exine formation. Immunofluorescence staining with JIM5 (pectin-specific) and LM10 (Xylan-specific) antibodies to detect pectin and Xylan levels in anthers of WT and *Zmsweet6a/6b* double mutant plants during and around pollen exine formation (stages S5 to S10). Green, signals from indicated anti-carbohydrate antibodies; blue, Calcofluor White signal. Signal in PE is indicated by white arrowheads. The imaging process for both mutant and WT specimens employed identical exposure durations. Bar, 20 μm.

We next examined xylan, another essential primexine scaffold component, using the LM10 antibody that specifically binds to the xylan backbone ^[34]^. In WT anthers at stage S6, LM10 signal localized clearly to the microspore surface, with a strong concurrent signal in metabolically active tapetal cells—suggesting synchronous xylan synthesis in the tapetum and deposition on the microspore surface (Fig. 4). In the double mutant, however, the LM10 signal in the tapetum was noticeably weaker; more critically, almost no signal was detected on the microspore surface (Fig. 4). By stages S9–S10, residual weak LM10 signal in the double mutant was confined to degenerating tapetal cells and failed to localize to the microspore surface, confirming a complete failure in xylan scaffold construction.

Together, these findings demonstrate that primexine deficiency in *Zmsweet6a/6b* results from a failure to assemble cohesive pectin and xylan scaffolds. This underscores the essential role of ZmSWEET6a/6b in providing the carbohydrate precursors required for the construction of the pollen wall template.

### 3.4. ZmSWEET6a and ZmSWEET6b function as plasma membrane hexose transporters to maintain sugar homeostasis in anther

Phylogenetic analysis places ZmSWEET6a and ZmSWEET6b within Clade II of the SWEET protein family (Fig. S1), whose members are typically localized to the plasma membrane and function primarily in hexose transport. Consistent with this classification, transient expression assays in *Nicotiana benthamiana* leaves and maize protoplasts confirmed that both ZmSWEET6a and ZmSWEET6b are specifically targeted to the plasma membrane (Fig. 5A, B). To assess their transport activity, we performed heterologous complementation in the hexose uptake–deficient yeast strain *EBY*.*VW4000. In vivo* transport assays demonstrated that expression of either ZmSWEET6a or ZmSWEET6b restored efficient uptake of glucose and fructose, confirming their identity as bona fide hexose transporters (Fig. 5C, D).

**Fig 5.**
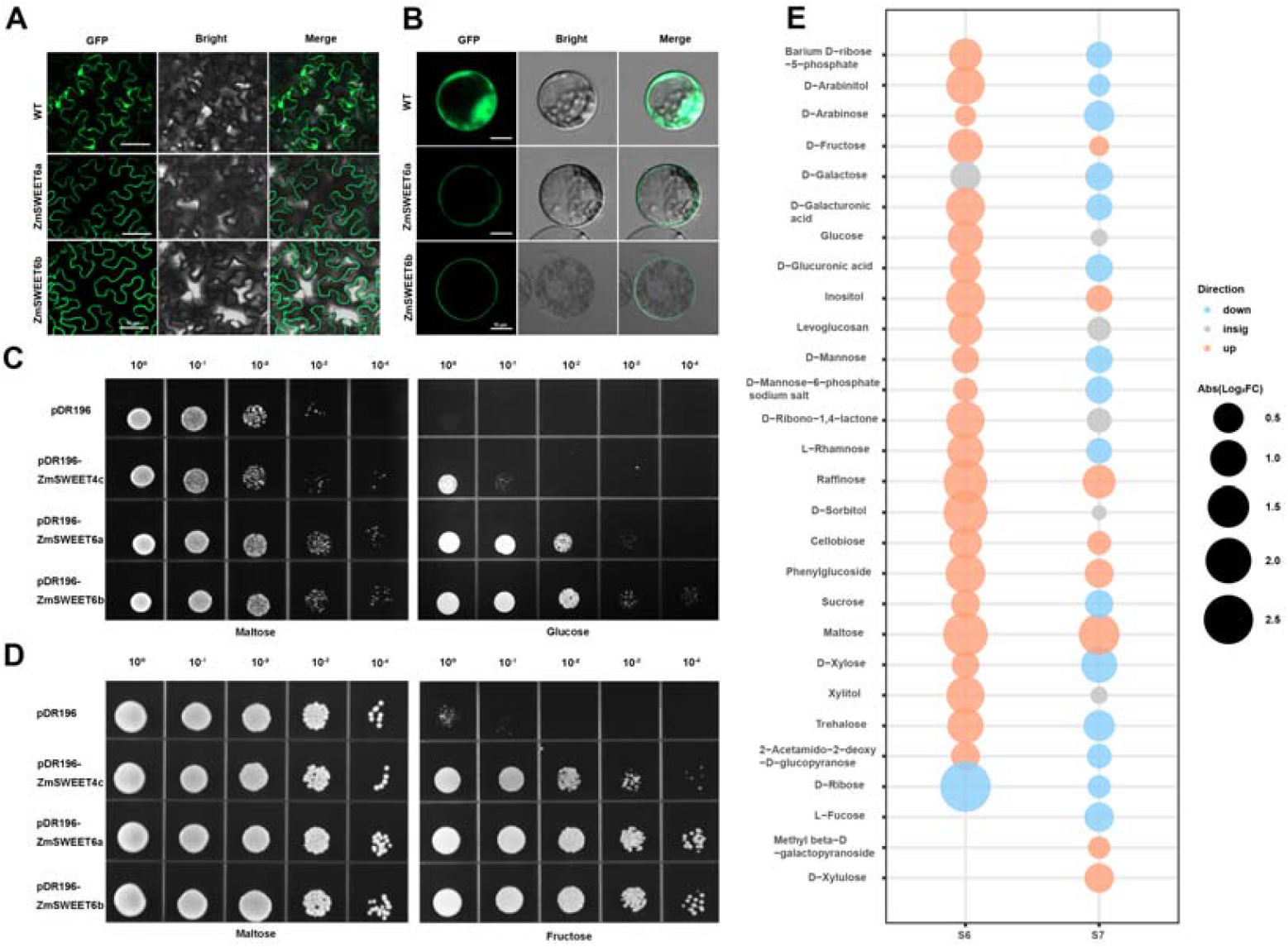
Loss of membrane-localized hexose transporters ZmSWEET6a and ZmSWEET6b disrupts anther sugar homeostasis. **(A-B)** Subcellular localization images of ZmSWEET6a and ZmSWEET6b, A was analyzed in *Nicotiana benthamiana* leaves, and B in protoplasts of maize Inbred line B73. The bar is as indicated in the Fig.. **(C-D)** Functional validation of the carbohydrate transport function of ZmSWEET6a and ZmSWEET6b. C and D show the verification of glucose and fructose transport capacities of ZmSWEET6a and ZmSWEET6b using the hexose-deficient yeast strain *EBY*.*VW4000*, respectively. **(E)** Targeted detection of monosaccharide, disaccharide, and trisaccharide contents in anthers of WT (LH244) and *Zmsweet6a/6b* double mutant plants using a GC-MS-based glycomics approach.

In flowering plants, phloem-derived sucrose is distributed throughout the anther via symplastic and apoplastic pathways ^[21]^. During this process, sugar transporters, including SWEETs, coordinate essential apoplastic fluxes that provide nutrients to developing microspores. During TEM analysis, we observed a striking phenotype in the *Zmsweet6a/6b* double mutant: massive starch accumulation in the endothecium of stage 6 anthers (Fig. S6A–C). This prompted us to investigate whether loss of ZmSWEET6a/6b disrupts sugar metabolism, potentially contributing to male sterility. We therefore performed comparative metabolite profiling of WT and mutant anthers at two key developmental windows, stages 6 and stages 7, using gas chromatography–mass spectrometry (GC-MS). Remarkably, sugar homeostasis was already severely perturbed by stages 6 in the double mutant. Levels of glucose, fructose, sucrose, sugar alcohols, and phosphorylated sugars were all significantly elevated compared to wild type (Fig. 5E, S6D–F; Table S2), indicating a systemic metabolic block. We hypothesize that the excess soluble sugar is channeled into starch as a transient storage strategy to relieve osmotic and metabolic stress, accounting for the prominent starch granules observed by TEM.

Interestingly, this compensatory response appears unsustainable. By stages 7, mutant anthers paradoxically exhibited reduced levels of soluble sugars, including sucrose, compared with the wild type (Fig. 5E, S6D–F; Table S2). This reduction may reflect feedback inhibition of sugar import or transport—a distinct form of sugar dysregulation indicative of a collapse in metabolic homeostasis. Notably, the precursors for xylan biosynthesis (D-xylose) and pectin biosynthesis (D-galacturonic acid) were also significantly reduced in *Zmsweet6a/6b* anthers at stages 7, consistent with our immunostaining results using LM10 and JIM5 antibodies that revealed impaired deposition of xylan and pectin (Fig. 4), respectively.

Loss of *ZmSWEET6a/6b* also triggers widespread transcriptomic reprogramming. RNA-seq analysis of stage S6 anthers identified 2,749 differentially expressed genes (DEGs) between WT and *Zmsweet6a/6b*, including 2,182 upregulated and 567 downregulated genes (Fig. S7A; Table S4). Functional enrichment analysis highlighted that DEGs were significantly enriched in pathways involved in polysaccharide breakdown and disaccharide biosynthesis. (Fig. S7B; Table S5). Consistent with the central role of sugar homeostasis in anther development, we observed dynamic transcriptional changes in sugar metabolism-related genes. Specifically, five *SUGAR TRANSPORT PROTEIN (STP)* genes were upregulated, whereas within the *SWEET* family, *ZmSWEET6a* and *ZmSWEET6b* were downregulated, while *ZmSWEET1b, 13a, 13b*, and *16* exhibited significant upregulation (Table S6). Trehalose-6-phosphate (Tre6P), a key sugar-signaling molecule ^[46]^ exhibited altered regulatory patterns, with three *TPS* genes (*ZmTPS10, 11, 13*) upregulated alongside *ZmTPS8* downregulation, while all five *TPP* genes (*ZmTPP1, 8, 10, 11, 12*) were upregulated. Additionally, invertase genes displayed changing in expression profiles, where *INVAN5* and *INCW8* were downregulated but *INCW1, IVR2*, and *IVR4* were upregulated. Moreover, three hexokinase genes (*HEX1, HEX4*, and *HEX7*) were all upregulated (Fig. S7C; Table S6). The differential expression of these genes likely reflectsa response to altered sugar homeostasis in the anthers.

Therefore, the metabolic changes and these transcriptional shifts—spanning transporters, signaling enzymes, and metabolic regulators—collectively suggest a systemic reconfiguration of sugar homeostasis in *Zmsweet6a/6b* anthers, likely representing adaptive responses to disrupted sugar allocation caused by ZmSWEET6a/6b loss, underscoring its pivotal role in coordinating carbohydrate dynamics during anther development.

### 3.5. Premature ROS burst triggers early programmed cell death in *Zmsweet6a/6b* anthers

Remarkably, DEGs between WT and *Zmsweet6a/6b* were significantly enriched in the “response to oxidative stress” category. Among the 52 DEGs in this category, 40—primarily encoding peroxidases or related proteins—were upregulated (Table S6), indicating that *ZmSWEET6a/6b* loss could perturb redox homeostasis. To address this, we stained anthers from both WT and mutant plants with H□DCF-DA. H□DCF-DA is a cell-permeable probe that, after intracellular deacetylation and ROS-mediated oxidation, generates green-fluorescent DCF for ROS detection and quantification ^[37]^. In the wild type, green fluorescence signal was weak in anthers from stage 6 to stage 10, with strong green fluorescence only becoming apparent in anthers at stage 11 and beyond. In contrast, in the *Zmsweet6a/6b* mutant, intense green fluorescence was already detectable as early as stage 6 and persisted throughout later stages of anther development (Fig. 6A, B). Furthermore, we also used Nitroblue tetrazolium (NBT) staining to detect superoxide anion (O□□), the major ROS component, in WT and *Zmsweet6a/6b* mutant anthers (Fig. 6C, D). Likewise, the *Zmswe*et6a/6b mutant anthers exhibited darker staining compared to the wild type, and cross-sections of stained anthers further revealed greater accumulation of superoxide (O□□) within the mutant (Fig. 6C, D). In that, both H□DCF-DA and NBT staining showed that *Zmsweet6a/6b* anthers accumulate higher levels of reactive oxygen species (ROS), with a ROS burst occurring as early as stage 6. Consistent with the observed accumulation of reactive oxygen species (ROS) in the mutant, RT-qPCR analysis revealed upregulated expression in *Zmsweet6a/6b* of four ROS scavenging–related genes (*ZmPODa, ZmMT2c, ZmTRXb*, and *ZmTrx*), two alternative oxidase family genes (*ZmAOX1* and *ZmAOX2*), and *ZmRBOH2*, which encodes a plasma membrane–localized NADPH oxidase involving in O□□ generation (Fig. S7D). In that, excessive ROS and O□L accumulation in *Zmsweet6a/6b* anthers might induce compensatory upregulation of scavenging genes, likely to mitigate oxidative stress.

**Fig 6.**
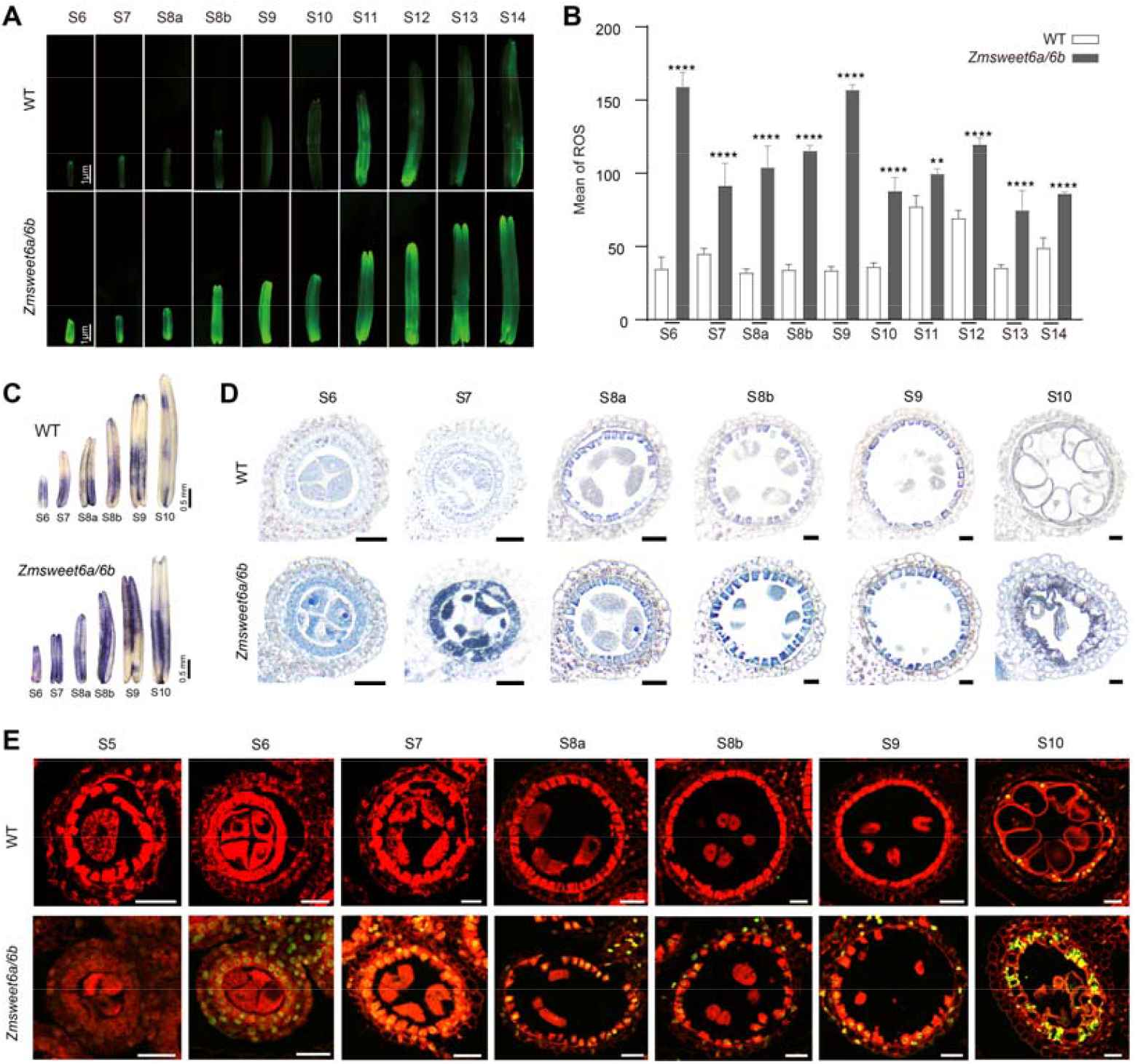
The loss of *ZmSWEET6a* and *ZmSWEET6b* triggers a burst of reactive oxygen species (ROS) and the premature onset of programmed cell death (PCD) in anthers. **(A-B)** Differences in reactive oxygen species (ROS) levels between WT and *Zmsweet6a/6b* double mutant anthers from S6 to S14, detected via H□DCF-DA staining. A, bar =1 nm. B, ** P < 0.01, **** P < 0.0001, Student’s *t*-test, n =3 biological replicates. **(C-D)** NBT staining of transverse sections of anthers from WT and *Zmsweet6a/6b* double mutant anthers from S6 to S10. C, bar=0.5 mm, D, bar=20 μm. **(E)** Programmed cell death (PCD) status of tapetal cells in transverse sections of WT and *Zmsweet6a/6b* double mutant anthers from S5 to S10, analyzed by TUNEL assay. Bar=25 μm. In A-E, each experiment was independently repeated three times.

Reactive oxygen species (ROS) burst is a hallmark signal of programmed cell death (PCD) ^[47, 48]^. We examined programmed cell death (PCD) in anthers of WT and *Zmsweet6a/6b* at different developmental stages using the terminal deoxynucleotidyl transferase-mediated dUTP nick end labeling (TUNEL) assay. Notably, TUNEL signals were observed in *Zmsweet6a/6b* anthers at stage 6, well before their appearance in WT anthers at stage 10 (Fig. 6C). Moreover, in WT anthers, normal PCD occurs exclusively in the tapetum, whereas in *Zmsweet6a/6b*, PCD is observed across all four anther wall cell layers. Remarkably, tapetal cell vacuolization in Zmsweet6a/6b anthers is initiated prematurely at stage 6, which may underlie the early initiation of PCD.

Together, these findings indicate that ZmSWEET6a/6b loss perturbs redox homeostasis, leading to excessive ROS accumulation and the premature, mislocalized initiation of PCD, which likely drives the defective anther development and male sterility in the mutant.

## 4. Discussion

### 4.1. ZmSWEET6a/6b-mediated hexose transport is essential for anther sugar homeostasis and early microsporogenesis

Anthers function as one of the primary sinks, consuming photosynthates during pollen development, with sugar transporters serving as indispensable mediators of this critical carbohydrate allocation that directly determines male fertility ^[49]^. This study demonstrates that the hexose transporters ZmSWEET6a and ZmSWEET6b act redundantly to support maize pollen development. In anther at meiotic stage, *ZmSWEET6a* and *ZmSWEET6b* are markedly higher expression level than those of other SWEET family genes and monosaccharide transporter-encoding genes (Table S1). Moreover, the mRNA *in situ* results suggested *ZmSWEET6a/6b* are expressed in the tapetum and PMCs (Fig. 1E), which suggests they play a major role in the asymptotically allocate hexose from anther wall to the meiocytes to supply the carbohydrate needs for gamete development. Thus, disruption of sugar transport in the *Zmsweet6a/6b* double mutant generates a ‘sink blockade,’ leading to hexose and sucrose accumulation that is ultimately converted into starch granules in the endothecium at stage 6, a layer that typically does not serve as a long-term starch reservoir (Fig. 3B, S5). Concurrently, five STP genes (*ZmSTP2, 3, 7, 14, 17*) and four *SWEET* genes are upregulated (Fig. S7C; Table S6), yet this compensatory response is insufficient to rescue the loss of ZmSWEET6a/6b function. This phenotype resembles that of the rice *csa* mutant, in which impaired sugar partitioning leads to excessive accumulation of sugars in leaves and stems, while floral organs exhibit reduced levels of both sugars and starch, ultimately causing carbon starvation and male sterility ^[27]^. Beyond sugar transporters, ZmSWEET6a/6b loss resulted in differential expression of a range of sugar metabolism and signaling genes, with a notable impact on those involved in Tre6P metabolism. Tre6P acts as a critical signal of sucrose availability in plants, forming a feedback regulatory loop: overexpression of *TPP* (trehalose-6-phosphate phosphatase) accelerates Tre6P degradation, thereby alleviating its repression on sucrose synthesis; the resulting increase in sucrose levels, in turn, enhances the activity of TPS (trehalose-6-phosphate synthase) ^[50]^. In *Zmsweet6a/6b* anthers, the upregulation of multiple *TPS* and *TPP* genes likely reflects a feedback response to the abnormal accumulation of sucrose (Fig. S7C; Table S6). Besides, the upregulation of *HEX1, HEX4*, and *HEX7* is consistent with the elevated glucose levels observed in the mutant anthers (Fig. S7C; Table S6). Overall, ZmSWEET6a/6b loss disrupts sugar composition in the anther and broadly affects sugar-related gene expression, emphasizing its critical role in sustaining sugar homeostasis during early microsporogenesis.

Notably, the transcriptomic and metabolomic analyses in this study were performed using whole anthers, thereby obscuring potential cell-type–specific regulatory effects. Future application of single-cell and spatially resolved transcriptomic approaches will be essential to resolve sugar-dependent regulatory networks within distinct anther cell types, particularly the tapetum, and to determine how ZmSWEET6a/6b-mediated carbon allocation modulates ROS homeostasis, tapetal programmed cell death, and ultimately pollen fertility.

### 4.2. Disrupted sugar transport triggers ROS burst and ectopic PCD, highlighting a conserved carbon–redox regulatory axis in anther development

Plant sugars, including disaccharides, raffinose family oligosaccharides, and fructans, function as direct reactive oxygen species (ROS) scavengers, particularly for hydroxyl radicals, thereby playing a critical role in maintaining ROS homeostasis ^[51]^. Here, the disruption of ZmSWEET6a/6b-mediated hexose transport in maize anthers leads to a cascade of metabolic disturbances that ultimately triggers an excessive accumulation of reactive oxygen species (ROS) as early as stage S6. The ROS burst is evidenced by intense green fluorescence in H□DCF-DA staining and dark staining in NBT assays, indicating significant superoxide accumulation (Fig. 6A-D). This premature ROS accumulation is directly linked to the early initiation of programmed cell death (PCD), as demonstrated by TUNEL assays showing PCD occurring as early as stage S6 in the mutant, compared to stage S10 in WT anthers (Fig. 6C). Notably, the PCD in *Zmsweet6a/6b* mutants is not restricted to the tapetum but occurs across all four anther wall cell layers, indicating a spatially aberrant cell death response (Fig. 6E). Convergently, multiple genetic models highlight the interplay between disrupted carbon metabolism and ROS-driven PCD in anther development. *ZmSWEET6a/6b* deficiency and CMS-C (*atp6c*) independently trigger ROS-mediated premature tapetal PCD: *ZmSWEET6a/6b* loss disrupts hexose transport, inducing metabolic stress and *HEX* gene upregulation (Table S6), while CMS-C impairs mitochondrial energy production ^[8]^. Similarly, maize *ms33* mutant exhibits ROS accumulation, early tapetal PCD, and energy starvation due to defective chloroplast lipid biosynthesis and starch turnover in the endothecium ^[7]^. The OsAGO2 pathway further reinforces this linkage by epigenetically suppressing *OsHXK1* to prevent ROS overproduction; its loss induces a ROS burst, culminating in male sterility ^[52]^. Notably, dysregulation of lipid metabolism represents another critical factor contributing to male sterility. In maize, disruption of the plastid-localized *ZmENR1/ZmHAD1* complex impairs anther cuticle and pollen wall formation, induces lipid peroxidation, and consequently triggers reactive oxygen species (ROS) accumulation and premature tapetal programmed cell death (PCD) ^[6]^. Although both lipid metabolism and the sugar metabolism pathway mediated by *ZmSWEET* genes fall under the broad category of carbon metabolism and ultimately converge on a conserved ROS-PCD cascade, their functional contributions to fertility are distinct. Sugar metabolism primarily supports energy provision and carbon skeleton allocation, thereby establishing the metabolic foundation for pollen development and indirectly influencing pollen wall biosynthesis. In contrast, lipid metabolism is more directly involved in maintaining membrane integrity, facilitating pollen wall assembly, and mediating lipid-based signaling, all of which are crucial for structural stability and coordinated development. Thus, through complementary physiological mechanisms, sugar and lipid metabolic pathways collectively fine-tune ROS homeostasis and ensure the proper timing of PCD in the anther. Disruption in either pathway can lead to dysregulated ROS dynamics and compromised male fertility.

Together, these findings underscore that disruption of carbon homeostasis—arising from defects in sugar transport, mitochondrial function, chloroplast regulation, or lipid metabolic imbalance—activates a conserved ROS-mediated programmed cell death cascade that is essential for maintaining fertility in plant reproductive tissues.

### 4.3. ZmSWEET6a/6b may ensure male fertility via a dual role in metabolic–redox coordination and primexine biosynthesis

The premature ROS burst in *Zmsweet6a/6b* mutants triggers early tapetal vacuolization (stage S6 vs. S10 in wild type), disrupting the developmental coordination required for pollen wall formation (Fig. 3, 6). While this ROS-PCD cascade is shared with other male-sterile mutants like *Zmms33*, maize CMS-C (*atp6c*), and *OsHXK1-OE*, which also exhibit ROS accumulation and premature tapetal PCD, the *Zmsweet6a/6b* phenotype is uniquely severe. Unlike these mutants that retain partial exine formation (with residual primexine-like structures and sporopollenin deposition) ^[8, 52, 53]^, *Zmsweet6a/6b* completely abolishes primexine assembly due to the absence of xylan and pectin accumulation as early as the pollen mother cell (PMC) stage (Fig. 4). This phenotype is consistent with that of the *Arabidopsis rpg1* (*AtSWEET8*) mutant, in which impaired hexose transport compromises primexine formation (Sun et al., 2013), but differs markedly in severity. Whereas *rpg1* exhibits partial primexine disruption with residual xylan and pectin deposition, loss of *ZmSWEET6a/6b* abolishes the initiation of primexine biosynthesis, leading to a complete failure of pollen wall formation (Fig. 3). Similarly, *Atrpg1* retains normal xylan/pectin composition but exhibits reduced sporopollenin deposition ^[15]^, contrasting with *Zmsweet6a/6b*’s complete absence of primexine scaffolding. In conclusion, our study identifies ZmSWEET6a/6b as a central integrator of carbon flux and redox signaling during maize pollen development. By sustaining sugar homeostasis, *ZmSWEET6a/6b* ensures proper ROS dynamics and the timely execution of tapetal programmed cell death, while concurrently supplying the hexose precursors required for primexine matrix assembly. These coordinated activities establish sugar transport as a key regulatory nexus coupling metabolic homeostasis to pollen wall biogenesis and male fertility, providing a conceptual framework for understanding how carbohydrate partitioning governs reproductive success in flowering plants, as well as offering potential genetic resources for crop hybrid breeding.

## Acknowledgements

This study was supported by the Biological Breeding-Major Projects (2023ZD04076), Natural Science Foundation of Tianjin (23JCZDJC00380) and Key Research and Development Program of Tianjin (25YFXNSN00030). Thanks to Prof. Xuexian Li and Prof. Xiaolei Sui from China Agricultural University for providing *EBY*.*VW4000* yeast strain.

## Author contributions

W.H and W.J. conceived this research, Y.Z., S.B. and F.S. conducted the experiments, analyzed the data, J.B. and Y.W. analyzed the transcriptome data. Z.D. provided guidance for the experiment and data analysis. Y.Z., S.B., W.H. wrote the manuscript and M.M. provided support for the writing. Z.D. and W.J. revised the manuscript. All authors have read and approved the contents of this paper.

