## Supplemental figure for "Functional Redundancy of ZmSWEET6a/b in Mediating Sugar Transport and Redox Homeostasis for Maize Primexine Formation"

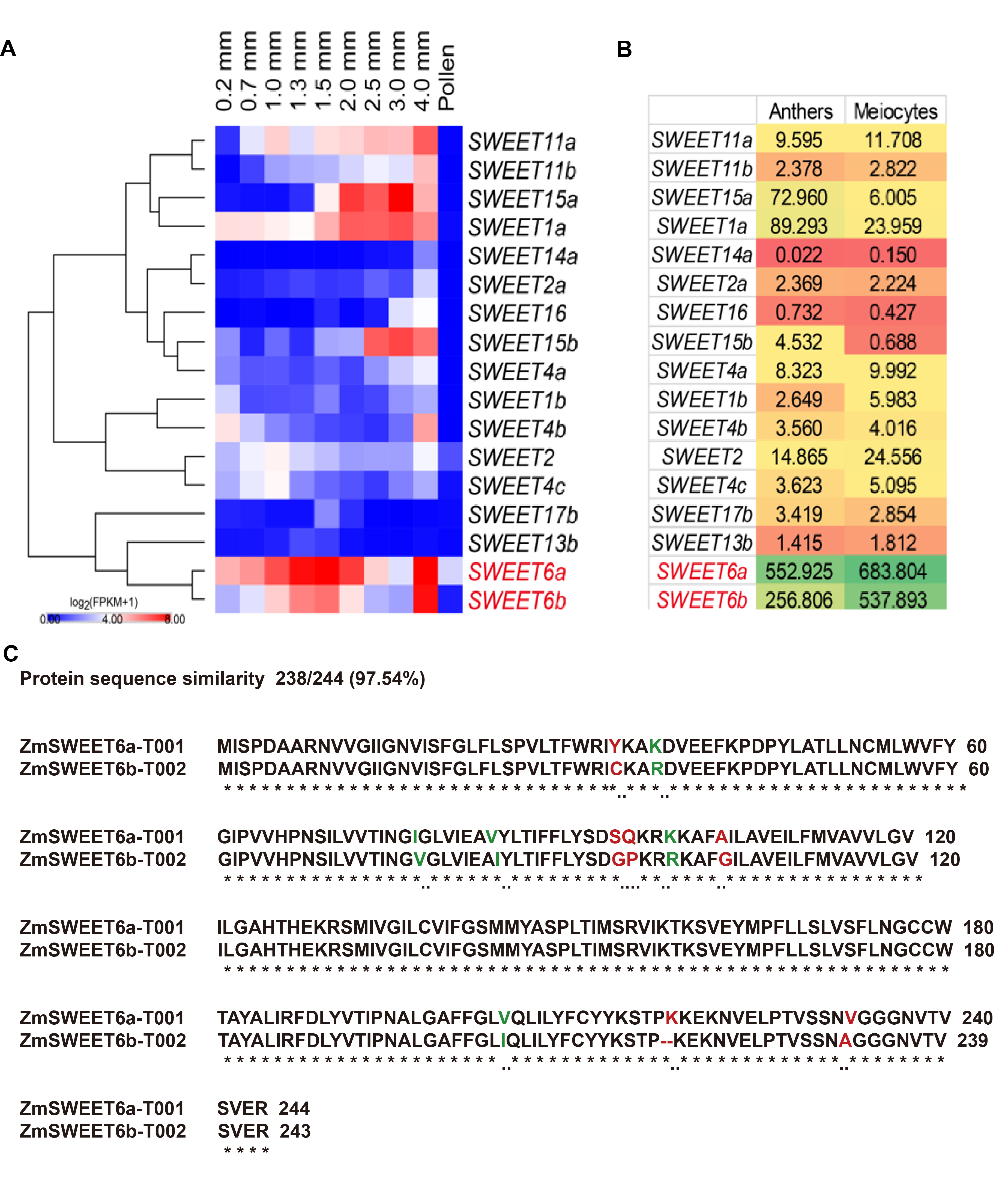

**Supplemental Fig. 1.** **Expression analysis of the SWEET family in maize. (A)** Expression profiles of ZmSWEET family genes in maize anthers and pollen grains of different lengths**. (B)** Expression profiles of ZmSWEET family genes in maize anthers and meiocytes**. (C)** The protein sequence similarity between ZmSWEET6a and ZmSWEET6b is as high as 97.5%. Data for panels A and B were derived from public datasets.

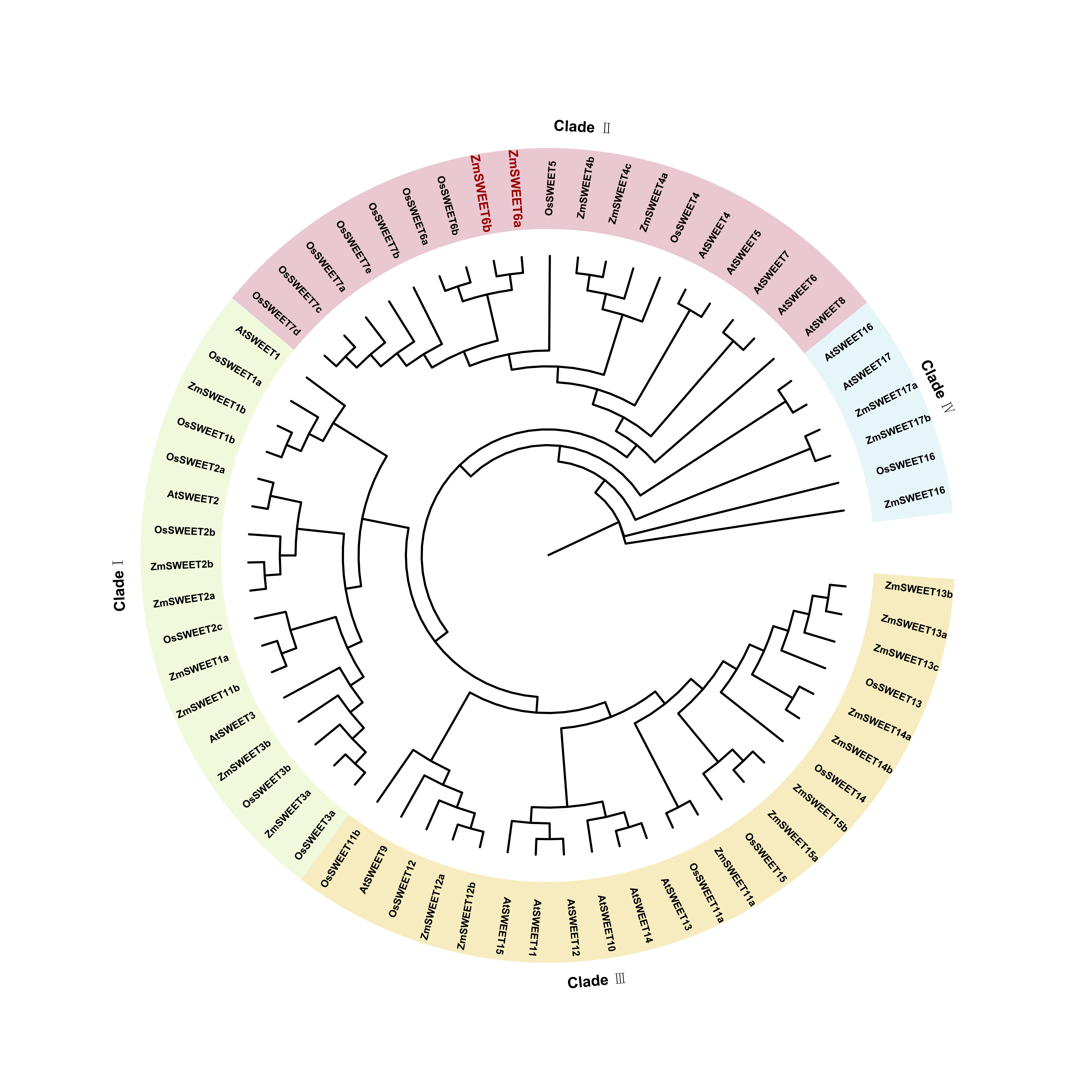

**Supplemental Fig. 2. Phylogenetic analysis was performed on SWEET family proteins from three plant species.** Proteins belonging to different clades are highlighted with colored backgrounds. Two ZmSWEET6 proteins from maize are marked in red.

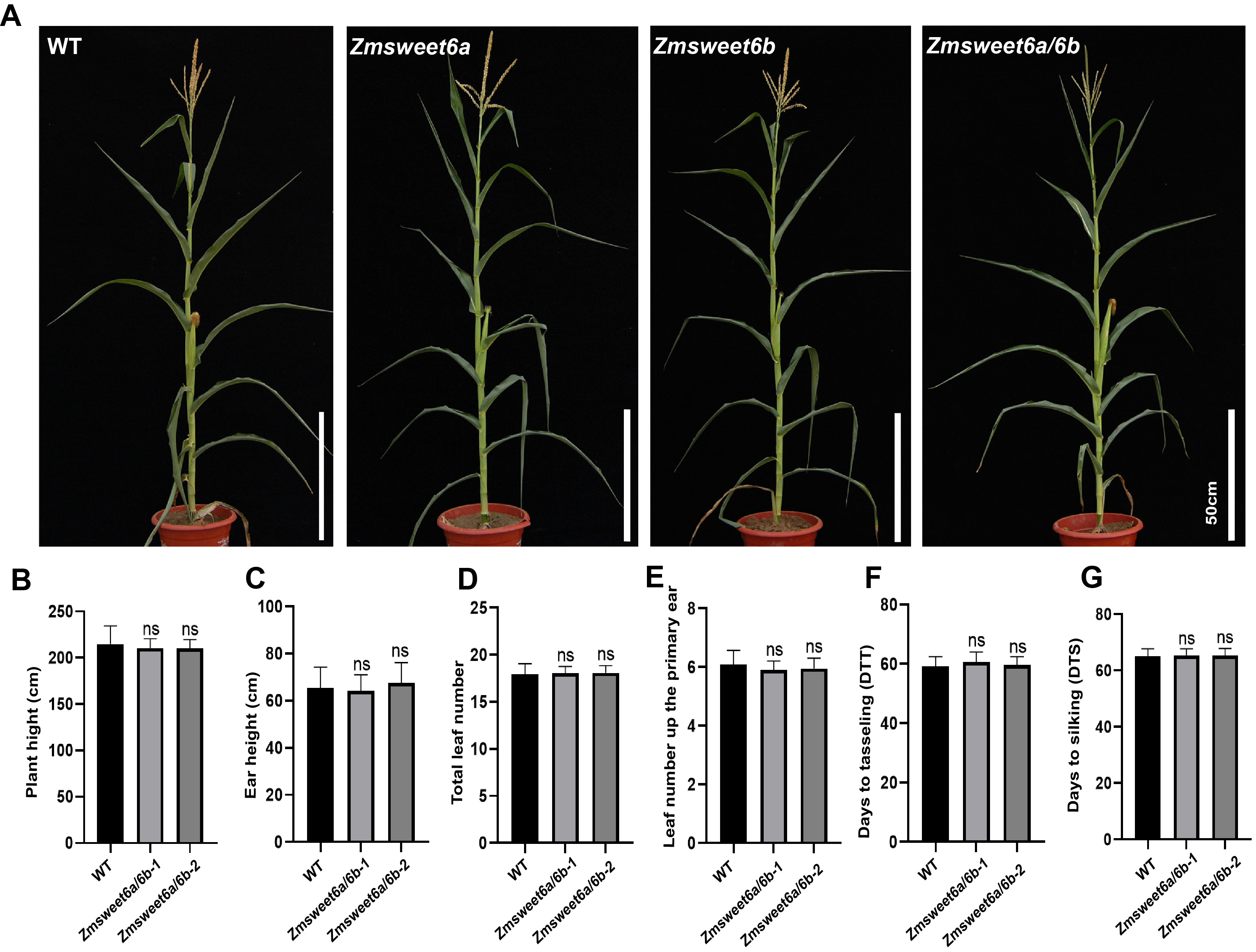

**Supplemental Fig. 3.** **Investigation of agronomic traits in WT and mutant Plants. (A)** Overall plant phenotype of the WT (LH244), *Zmsweet6a* single mutant, *Zmsweet6b* single mutant and *Zmsweet6a/6b* double mutant. Bar, 50 cm. **(B-G)** Phenotypic investigation of plant height and other agronomic traits at flowering stage in WT and two Independent *Zmsweet6a/6b* double mutant lines.

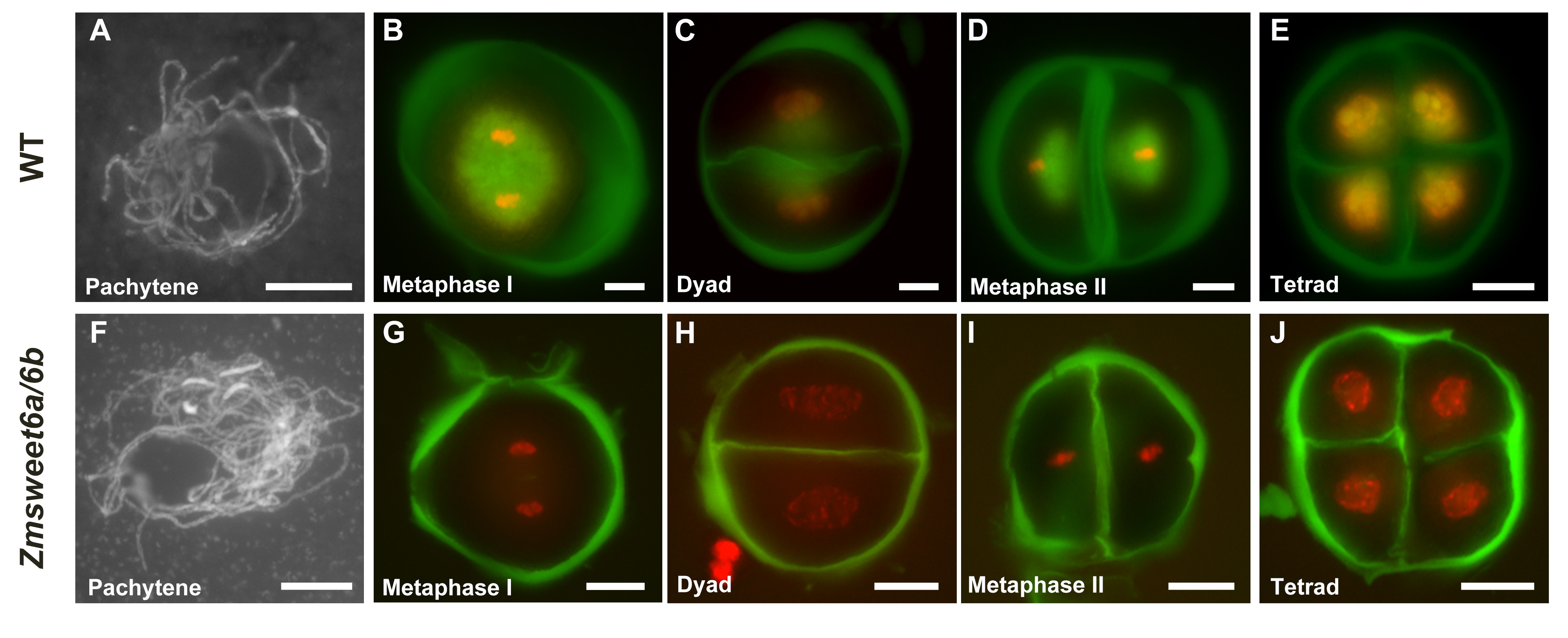

**Supplemental Fig. 4. The meiotic process and callose deposition exhibited no abnormalities between WT and *Zmsweet6a/6b* double mutant.** DAPI staining revealed chromosomal behavior in WT and *Zmsweet6a/6b* double mutants, while toluidine blue staining indicated callose deposition patterns. A and F display DAPI staining only; B - E and G - J, chromosomes are indicated in red and callose is labeled in green. The imaging process for both mutant and WT specimens employed identical exposure durations. Bar, 20 μm.

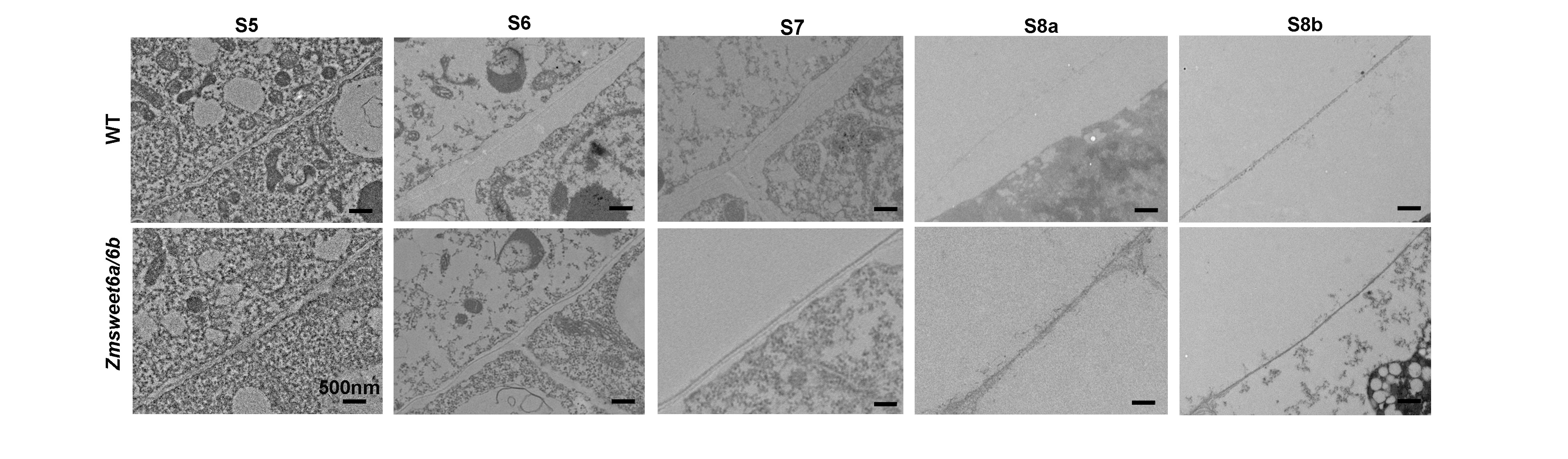

**Supplemental Fig. 5. No phenotypic differences in ubisch bodies were observed between the WT and *Zmsweet6a/6b* double mutant during the S5 and S8b**. Bar, 500 nm.

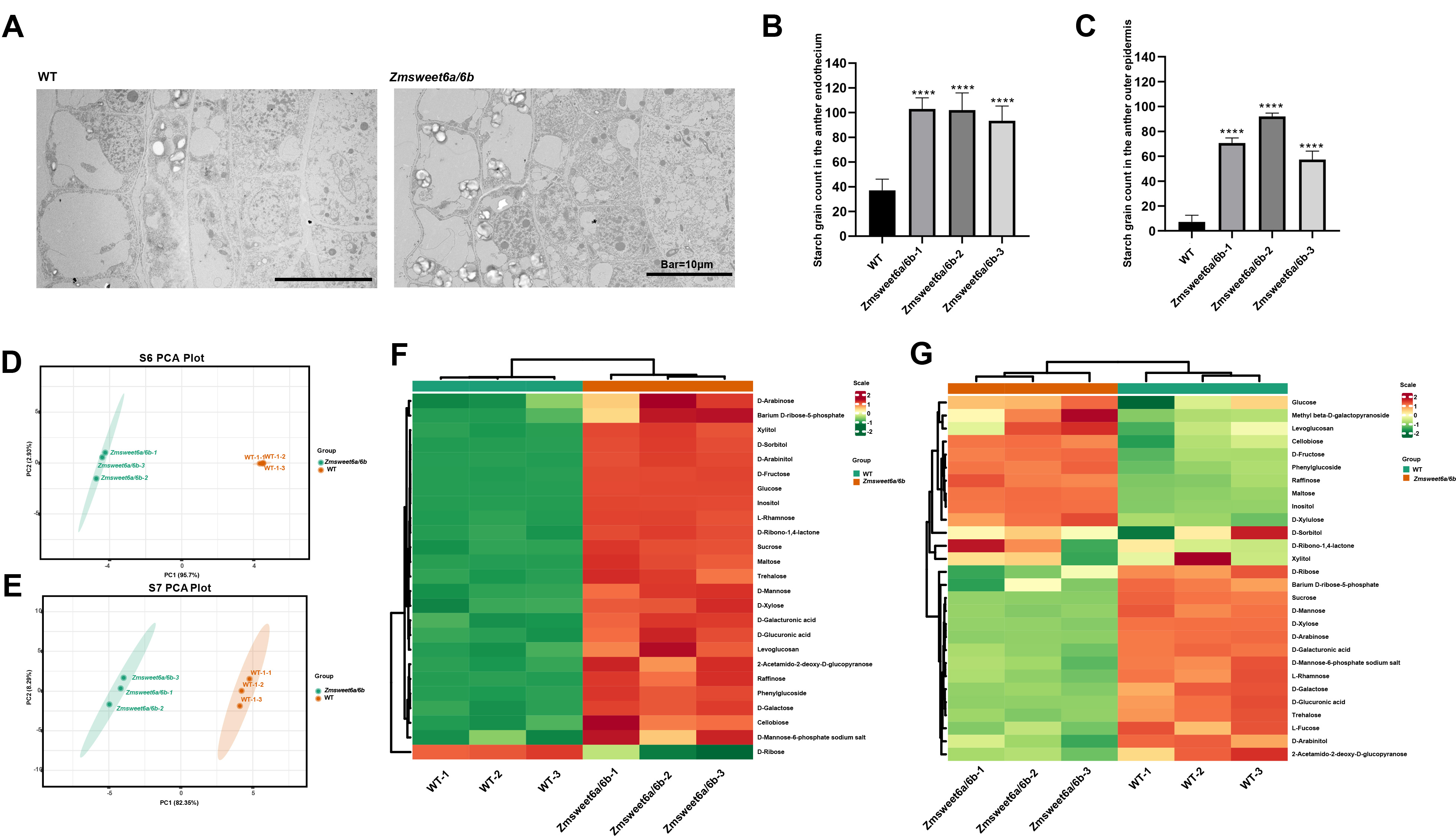

**Supplemental Fig. 6.** **The anthers of *Zmsweet6a/6b* double mutant exhibit sugar metabolism disorders during the stages 6-7, with a significant accumulation of starch grains observed specifically at the stage 6.** **(A)** Ultrastructural observation of anther cross-sections of WT and *Zmsweet6a/6b* double mutant at the stage 6 via transmission electron microscopy (TEM). **(B-C)** Quantitative analysis of starch grains in the endothecium (B) and outer epidermis (C) of anthers from WT and *Zmsweet6a/6b* double mutant. **** P < 0.0001, Student’s *t*-test, n =9 (3 anthers×3 cells/anther) **(D F****)** Principal Component Analysis (PCA) of anther metabolomics data from WT and *Zmsweet6a/6b* double mutant during the stage 6 (D) and stage 7 (F). **(E G)** Heatmap of cluster analysis for anther metabolomics data from WT and *Zmsweet6a/6b* double mutant during the stage 6 (E) and stage 7 (G). In D-G, n=3 biological replicates.

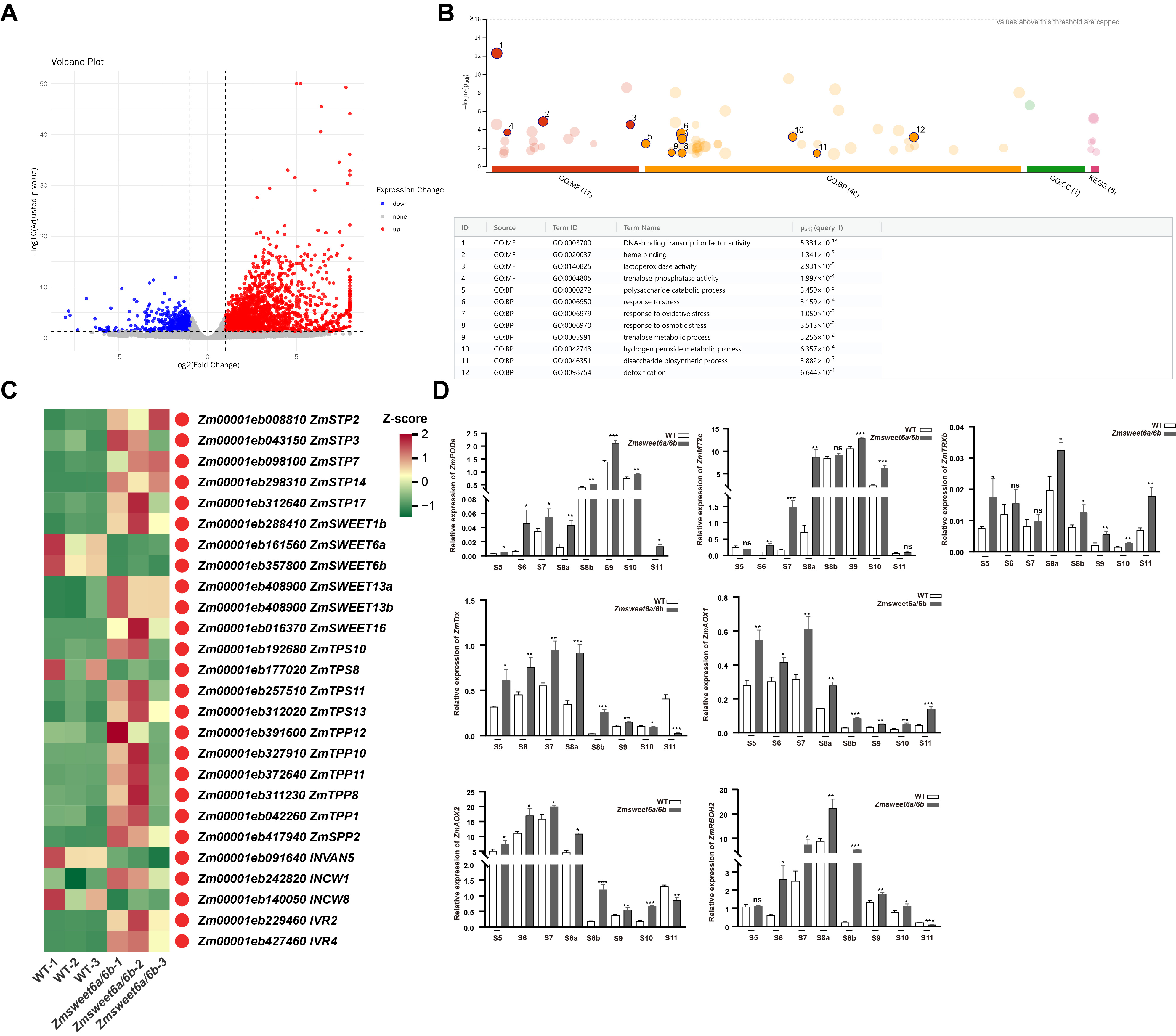

**Supplemental Fig. 7. Transcriptomic profiling of anthers from the WT and *Zmsweet6a/6b* double mutants, followed by qPCR validation of redox-associated genes. (A-B)** Volcano plot (A) and GO enrichment plot (B) of transcriptomes of anthers from WT and *Zmsweet6a/6b* double mutant plants at stage 6, with three biological replicates; each replicate contained a mixed sample of three or more independent lines. **(C)** Transcriptomic profiling of sugar transport-associated genes via RNA-Seq analysis, Q-value < 0.05, n =3 biological replicates. **(D)** Determination of expression pattern changes of *ZmPODa*, *ZmMT2c*, *ZmTRXb*, *ZmTrx*, *ZmAOX1*, *ZmAOX2*, and *ZmRBOH2* in WT and *Zmsweet6a/6b* double mutant from the S5 to S11 via quantitative real-time polymerase chain reaction (qPCR). * P < 0.01, ** P < 0.01, *** P < 0.001, Student’s *t*-test, n =3 biological replicates.
